# The interplay of HSP90s with YDA regulates main body axis formation during early embryogenesis in Arabidopsis

**DOI:** 10.1101/2020.09.29.319384

**Authors:** Despina Samakovli, Tereza Tichá, Tereza Vavrdová, Natálie Závorková, Ales Pecinka, Miroslav Ovečka, Jozef Šamaj

**Author notes:** These three authors contributed equally to this work (Despina Samakovli, Tereza Tichá, Tereza Vavrdová).

## Abstract

The YODA kinase (YDA) pathway is intimately associated with the control of *Arabidopsis thaliana* embryo development but little is known regarding its regulators. Using genetic analysis, HEAT SHOCK PROTEINS 90 (HSP90s) emerge as potent regulators of YDA in the process of embryo development and patterning. This study is focused on the characterization and quantification of early embryonal traits of single and double *hsp90* and *yda* mutants. The mutant analysis was supported by expression analyses of cell-specific *WUSCHEL-RELATED HOMEOBOX* 2 (*WOX2*) and *WOX8* genes during early embryonic development. Chromatin immunoprecipitation assays corroborated the involvement of YDA and HSP90s in the epigenetic control of chromatin remodeling during early embryogenesis. Genetic interactions among HSP90s and members of the YDA signaling pathway affected the development of both embryo proper and suspensor. Impaired function of HSP90s or YDA had an impact on the spatiotemporal expression of *WOX8* and *WOX2* suggesting their essential role in cell fate determination and interference with auxin distribution. Hence, the interplay between HSP90s and YDA signaling cascade mediates the epigenetic control regulating the transcriptional networks shaping early embryo development.

## Introduction

At the onset of embryogenesis the main body axis formation is the first critical step in the development of an organism. Axis formation in plants occurs during the elongation and subsequent asymmetric division of the zygote, establishing a shootward upper part and a rootward lower part. During the early embryonic development of Arabidopsis, the initiation of this apical-basal axis coincides with the longitudinal axis formed by unfertilized egg cell and zygote that have the nucleus at the apical half and vacuoles at the basal part (Mansfield and Briarty, 1991; Jürgens and Mayer, 1994). This early development follows a well-defined pattern in which a complex network of specific regulatory genes interact with cellular polarity pathways to mark the development of new domains through the regulation of cell division occurrence and symmetry. During its asymmetric division, the zygote partitions into a small apical daughter cell and a large vacuolated basal cell. The first one gives rise to the apical embryo lineage that will form the proembryo. The lower daughter cell will further develop into the suspensor that connects the embryo to maternal tissue (Mansfield and Briarty, 1991; Jürgens and Mayer, 1994). The uppermost cell of the suspensor cell lineage, called the hypophysis, is incorporated into the embryo and contributes to the formation of the root meristem (Dolan *et al*., 1993).

Two major pathways contribute to the establishment of apical-basal axis formation: the WUSCHEL-RELATED HOMEOBOX (WOX) transcription factors-dependent and the auxin-dependent (Lau *et al*., 2012). *WOX* gene family members are expressed in the zygote and considered as main regulators of early embryo development (Haecker *et al*., 2004; Wu *et al*., 2007; Breuninger *et al*., 2008; Ueda *et al*., 2011). *WOX2* is strictly expressed in the shootward embryonic lineage and controls gene expression programs required for the initiation of the shoot meristem. By controlling auxin to cytokinin ratios, it maintains the proliferation activity of stem cell progenitors (Zhang *et al*., 2017). *WOX8* and *WOX9* are expressed in the rootward embryonic lineage and regulate suspensor development. Their activity regulates WOX2 and auxin patterning in a non-cell-autonomous way during embryo development (Haecker *et al*., 2004; Wu *et al*., 2007; Breuninger *et al*., 2008). WRKY2 (WRKY DNA-BINDING PROTEIN 2)-dependent transcription of *WOX8* connects zygote polarization with the early patterning of the embryo, while WRKY2 is phosphorylated downstream of the YODA (YDA) signaling cascade (Ueda *et al*., 2017).

Mitogen-activated protein kinases (MPKs) are key regulators of conditional signaling in embryonic and post-embryonic development (reviewed in Komis *et al*., 2018). YDA pathway is one of the major signaling cascades controlling cell division plane (CDP) orientation and cell division symmetry. It confers transcriptional control during stomata ontogenesis and zygote development by targeting and phosphorylating transcription factors involved in specific developmental processes (Lukowitz *et al*., 2004; Bergmann and Sack, 2007; Dong *et al*., 2009; Jeong *et al*., 2011; Zhang *et al*., 2015). *YDA* encodes a MPK kinase kinase involved in zygote elongation and the shootward extra-embryonic cell fate (Lukowitz *et al*., 2004). Activation of downstream MPK kinases depends most likely on the paternally derived kinase SHORT SUSPENSOR (SSP; Bayer *et al*., 2009) while SSP paralogs, like BRASSINOSTEROID SIGNALING KINASE 1 and 2 (BSK1 and BSK2) participate in YDA signaling processes including embryogenesis (Neu *et al*., 2019). Zygote elongation and the initiation of the first asymmetric division relies on the nuclear protein GROUNDED (GRD; Jeong *et al*., 2011, Waki *et al*., 2011). Additional signaling components, like the peptide EMBRYO SURROUNDING FACTOR 1 (ESF1), produced in the central cell before fertilization, act together with SSP to control the elongation of the suspensor through YDA signaling cascade (Costa *et al*., 2014).

Epigenetic programs such as DNA methylation and histone modifications control gene expression patterns during embryo and seed development (Feng *et al*., 2010; Moazed, 2011). Polycomb group proteins (PcG) form Polycomb Repressive Complex 1 and 2 (PRC1 and PRC2), which regulate embryo development by suppressing the expression of specific genes (Schwartz and Pirrotta, 2013).

Recently, we showed that HEAT SHOCK PROTEINS 90 (HSP90s) interact with YDA signaling cascade to control stomata patterning and distribution by modulating MPK6 phosphorylation and the transcriptional activation of SPCH both under control and heat stress conditions (Samakovli *et al*., 2020b). HSP90s are evolutionarily conserved molecular chaperones involved in the regulation of a plethora of signaling proteins including kinases and transcription factors (Wandinger *et al*., 2008; Neckers *et al*., 2009; Taipale *et al*., 2010). HSP90s are essential in many processes encompassing development, homeostasis and organismal evolution (Rutherford and Lindquist, 1998; Queitsch *et al*., 2002; Samakovli *et al*., 2007; Jarosz *et al*., 2010; Margaritopoulou *et al*., 2016). Expression of *HSP90s* is tightly regulated during embryo development showing dramatic upregulation of *HSP90.1* in developing embryos, induction of *HSP90.2* expression during pod elongation and a slight decrease in mature embryos (Prasinos *et al*., 2005), while *hsp90* mutants display lethal defects in embryogenesis (Samakovli *et al*., 2007). Studies showed interactions between HSP90s and histones (Schnaider *et al*., 1999) and the positive role of HSP90s in chromatin condensation leading to decreased dissociation rates of histones (Csermely *et al*., 1994). Additionally, HSP90s directly interact with Trithorax (Trx), suggesting a role of HSP90s in the maintenance of active expression state of target genes (Tariq *et al*., 2009).

In this study, we demonstrate the relationship between HSP90s and YDA signaling pathway during Arabidopsis embryo development. We show that impaired function of HSP90s and/or YDA signaling cascade leads to defective development of the suspensor, aberrant spatiotemporal transcriptional activation of *WOX8* and *WOX2* genes and deregulated auxin maxima in developing embryos. Moreover, HSP90s are involved in the epigenetic control of *WOX8, WOX9* and *WOX2* genes by controlling the occupancy of Histone 3 (H3) at the WRKY binding sites. A negative correlation between HSP90s and YDA activity in the distribution of nucleosomes at WRKY binding sites residing in the promoters of *WOX8*, *WOX9* and *WOX2* was revealed. Our results suggest that HSP90s and YDA together contribute to the correct basal and apical cell fate initiation and suspensor elongation by modulating the transcriptional networks. Thus, HSP90s are new players involved in the main body axis formation during early embryogenesis in Arabidopsis.

## Materials and Methods

### Plant material and growth conditions

Seeds of *Arabidopsis thaliana* (L.) Heynh were grown under axenic conditions on Phytagel (Sigma, Czech Republic) solidified half-strength Murashige Skoog medium supplemented with 1% w/v sucrose, as previously described (Beck *et al*., 2010). Columbia (Col-0) and Landsberg *erecta* (L*er*) ecotype were used as wild-type. The mutant genotypes of interest used in this study are summarized in Table S1. For the experiments were used *proDR5:GFP* (Friml *et al*., 2003), *proWOX8:NLS-YFP* and *proWOX2:NLS-YFP* lines (Breuninger *et al*., 2008) and their crosses with mutant lines; and the lines *proHSP90.1:GUS* (Haralampidis *et al*., 2002), *proHSP90.2:GUS* (Prasinos *et al*., 2005), *proYDA:GUS* (Meng *et al*., 2016) and *proHSP90.1:HSP90.1-GFP* (Samakovli *et al*., 2014), *proHSP90.2:HSP90.2-GFP* (Samakovli *et al*., 2014), *proYDA:YDA-YFP* (Zhang *et al*., 2015). The analysis of crosses was performed at F3 generation in homozygous mutants selected after genotyping unless stated otherwise. The mutants crossed with the lines *proWOX2:NLS-YFP* or *proWOX8:NLS-YFP* were selected by phosphinotricin. The primers used for genotyping are included in Table S2.

### Phenotypic and microscopic analyses

For Nomarski microscopy, siliques with immature seeds were fixed in ethanol with acetic acid (3:1 v/v), the extracted embryos from siliques were mounted in chloral hydrate solution (8 g chloral hydrate, 1 ml glycerol, and 3 ml water). Modified pseudo-Schiff propidium iodide staining was performed as previously described (Truenit *et al*., 2008). Histochemical β-glucuronidase (GUS) activity assays were performed on siliques as previously described (Samakovli *et al*., 2020b) and immature seeds were extracted after staining on slides. SCRI Renaissance 2200 staining was performed as previously described (Musielak *et al*., 2015).

When analysing embryos of reporter lines, seeds were extracted onto slide with water and immediately observed. GUS staining documentation and embryonal analysis of marker lines using Nomarski microscopy were performed on light microscope Zeiss Axio Imager M2 equipped with DIC optics (Zeiss, Germany) and Zeiss Zen 2012 Blue software (Zeiss, Germany). Samples stained with propidium iodide and embryos expressing *proHSP90.1:HSP90.1-GFP, proHSP90.2:HSP90.2-GFP*, or *proYDA:YDA-GFP* were observed on confocal laser scanning microscope (LSM 710, Zeiss, Germany) and analysed with Zeiss Zen 2012 Black software (Zeiss, Germany).

### Imaging and post-acquisition processing

Image acquisition was done by individual optical sections in the Z-stack mode. Thickness of optical sections was adjusted according to selected objective and excitation wavelength using the Nyquist sampling function of the ZEN software. The post-processing of Z-stacks from confocal laser scanning microscope was used for creation of maximum intensity projections from individual z-stacks. In images from epifluorescence microscope the medial optical sections were selected and representative subsets in total from three to five independent optical sections along the median plane were created, corresponding to a respective distribution of fluorescence intensity. The brightness and contrast of all optical sections in the subset were uniformly adjusted. Created maximum intensity projections were processed by the Nearest Neighbour or Constrained Iterative deconvolution algorithms with default settings. After uniform correction of the brightness and contrast, fluorescence intensity of *WOX8* expression levels in developing embryos from maximum intensity projections of subsets were semi-quantitatively evaluated using pseudo color-coded range distribution. The range from zero fluorescence intensity (dark blue) to maximum fluorescence intensity (white) was designed by arbitrary units in Zen 2012 Blue software. The images of propidium iodide stained embryos (originally showing red stained cell walls) were inverted to black-white contrasted mode in ImageJ (Schneider *et al*., 2012). Post-processing and editing of images were done using ZEN 2014 software (Carl Zeiss, Germany) and final figure plates were obtained using Photoshop 6.0/CS and Microsoft PowerPoint software.

### Calculation of Epistasis Value

To study the effect of epistasis, we used a formula that can show the strength and direction of the epistasis (Hartman *et al*., 2001; Segrè *et al*., 2005). The difference of observed double mutant phenotype versus the predicted double mutant phenotype was normalized, considering that there was additivity of the single mutants. The epistasis value was then normalized to the wild-type. The phenotype for the wild-type was set as w, mutant a as a, mutant b as b, and double mutant as ab. The epistasis value was calculated as {ab – [w + (a – w) + (b – w)]/w}. If the epistasis value was positive, it showed evidence for synergistic epistasis, while antagonistic epistasis was suggested by negative values. The larger the epistasis value, the stronger the epistatic effects.

### *In silico* and statistical analysis

For the prediction of WRKY binding sites on the promoters of genes of interest the AtcisDB – Arabidopsis cis-regulatory element database (https://agris-knowledgebase.org/AtcisDB/) was used. Descriptive data measured in Zeiss Zen 2012 Blue software (e.g. suspensor length, signal intensity), were subjected to statistical analysis. Generally, datasets were first subjected to Shapiro-Wilk test of normality and Levene’s test of homogeneity, then, either Welch’s one-way ANOVA, one-way ANOVA or Kruskal-Wallis tests were calculated. The ANOVA tests were followed by either Tukey’s honestly significant difference corrected for unequal sample size or Scheffé’s test. The χ^2^ tests were calculated according to Pearson’s formula.

### Chromatin immunoprecipitation (ChIP)

ChIP experiments were performed as described previously (Yamaguchi *et al*., 2014), in three biological replicates, using the following antibodies: polyclonal rabbit anti-HSP90 IgG (Santa Cruiz, USA), polyclonal rabbit anti-H3 (Abcam, UK). DNA recovered after ChIP and DNA from input chromatin was extracted by DNA isolation kit (Macherey-Nagel, Germany) and then analysed by qPCR. The qPCR was performed as described (Samakovli *et al*., 2020b) with primers listed in Table S2.

## Results

### HSP90 mutations partially rescue embryonic suspensor phenotypes in loss- and gain-of-function *yda* mutants

Previous reports showed that homozygous loss-of-function *yda-1* and gain-of-function *ΔNyda* mutants, harboring a constitutively activated form of YDA lacking the N-terminal regulatory domain, cannot enter the reproductive phase exhibiting sterile phenotypes (Meng *et al*., 2012). To investigate the genetic interactions of *HSP90s* and *YDA* genetic loci during embryo development, we used crosses of *yda-1* and *ΔNyda* mutants with *hsp90.1* and *hsp90.2* knockouts or *proRAC2:HSP90^RNAi^* line (*hsp90^RNAi^*) driven by *RAC2/ROP7* promoter that results in the silencing of all four HSP90 cytoplasmic members (Samakovli *et al*., 2020b). *RAC2* promoter has a moderate expression in developing siliques (Klepikova *et al*., 2016). The *hsp90^RNAi^* line was used to overcome the limitation that it is not possible to prepare *hsp90.1 hsp90.2* double mutants, as plants homozygous for *hsp90.1* and heterozygous for *hsp90.2* displayed reduced fecundity unable to produce offspring (Hubert *et al*., 2009). An epistatic relation between *HSP90.1* and *YDA* was previously reported (Samakovli *et al*., 2020b), as *hsp90.1* mutation suppressed the severe phenotypes of *yda-1* and *ΔNyda* resulting in fertile double mutant plants (Samakovli *et al*., 2020b). By contrast, *hsp90.2* could not restore *ΔNyda* phenotype as homozygous plants produced no inflorescence, while no homozygous *hsp90.2 yda-1* mutants from *hsp90.2 yda-1/+* segregating plants were received.

In agreement with published transcriptome analysis data (Hofmann *et al*., 2019), we showed expression of *HSP90.1, HSP90.2* and *YDA* in developing embryos from the globular up to torpedo stages by using transcriptional fusions of *proHSP90.1:GUS*, *proHSP90.2:GUS* and *proYDA:GUS* and translational fusions of *proHSP90.1:HSP90.1-GFP*, *proHSP90.2:HSP90.2-GFP* and *proYDA:YDA-GFP* expressing lines (Supplemental Fig.S1).

Genetic impairment of *YDA* or *MPK3* and *MPK6* restricts zygote elongation leading to the production of basal cell that has almost the same size as the apical one (Lukowitz *et al*., 2004; Wang *et al*., 2007). The following cell divisions of the basal and the apical cell are also aberrant resulting in embryos without or with hardly obvious suspensor. In contrast, the constitutive activation of YDA results in embryos with long suspensors (Lukowitz *et al*., 2004).

Since HSP90s are involved in the regulation of YDA signaling cascade during stomatal development (Samakovli *et al*., 2020b), a model of asymmetric cell lineage, we analyzed embryonic phenotypes of single and double *hsp90* and *yda* mutants. To this end, we analyzed and measured the suspensor and proembryo length at the early globular stage of single and double *hsp90* and *yda* mutants (Fig.1). The *yda-1* mutants showed very short suspensors contrary to very long ones in *ΔNyda* (Fig.1, A and B) corroborating previous studies (Lukowitz *et al*., 2004). The phenotypic analysis showed that in *hsp90.1* suspensor length was shorter than in the wild-type, whereas in *hsp90.2* and *hsp90^RNAi^* suspensor lengths were similar to the wild-type ones (Fig.1A and B). Interestingly, *hsp90.1* genetic depletion suppressed the suspensor phenotypes of both *yda-1* and *ΔNyda*, as in *hsp90.1 yda-1* and *hsp90.1 ΔNyda* the suspensor length was similar to the wild-type (Fig.1A and B). We observed a similar effect upon silencing of all cytoplasmic HSP90 members using *hsp90^RNAi^* line in the embryos of *yda-1* and *ΔNyda*, since the suspensor length of *hsp90^RNAi^ yda-1* and *hsp90^RNAi^ ΔNyda* was similar to the one of wild-type embryos (Fig.1A and B). In *hsp90.2 yda-1 and hsp90.2 ΔNyda* double mutants, the suspensor length was higher compared to the wild-type (Fig.1A and B).

**Fig.1.**
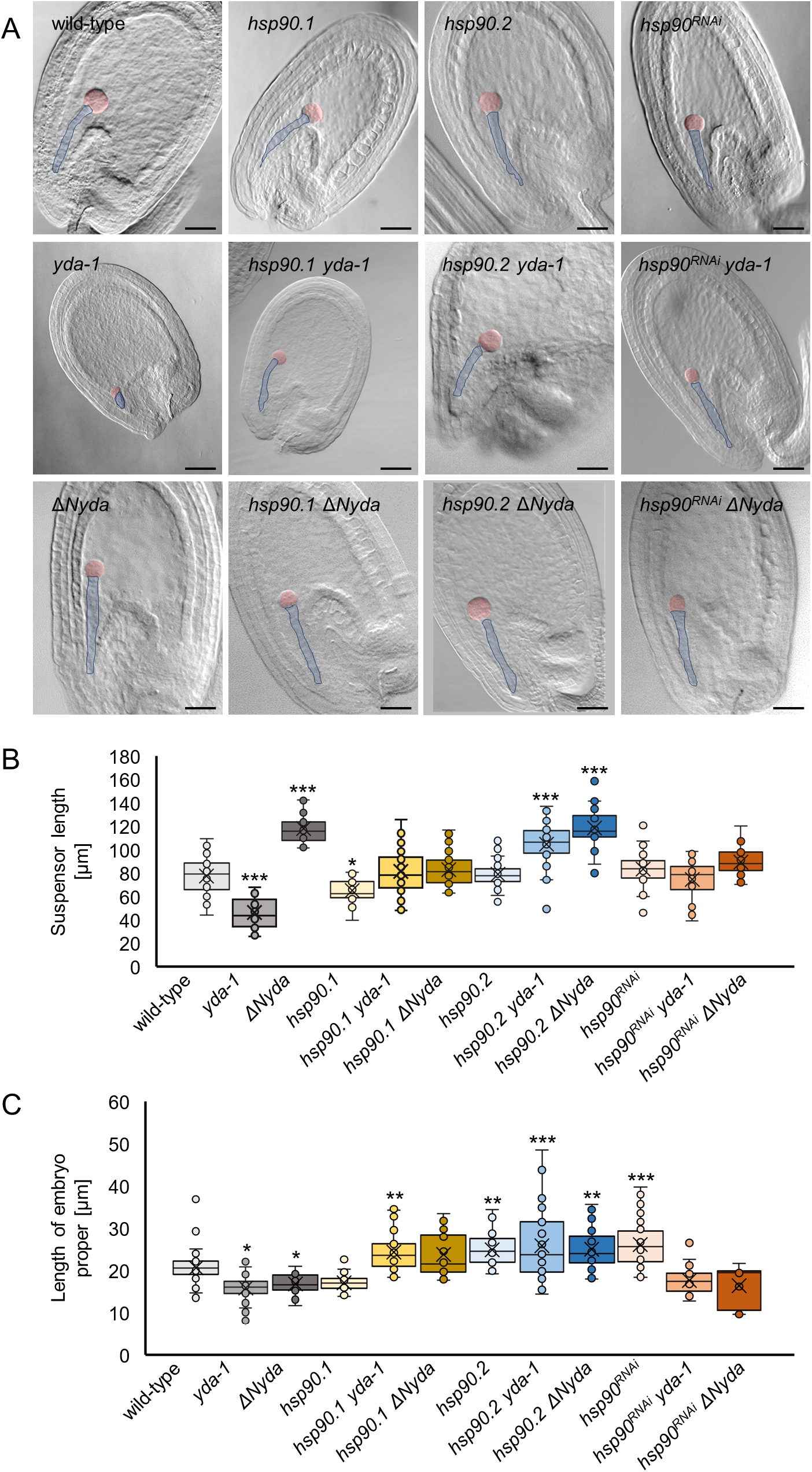
Overview of embryonic development in *hsp90, yda* single and double mutants. (A) DIC images of cleared whole-mount seeds containing embryos at the globular stage of the indicated genotypes. Suspensor is traced with the blue line, embryo proper is marked in red. Scale bars, 50 μm. (B) Quantitative analysis of the suspensor length in the indicated genotypes (N≥30) at the globular stage (Welch’s ANOVA followed by Tukey’s test, statistically significant differences compared to WT are shown, *** is significant at p <0.001, * is significant at p<0.05). (C) Quantitive analysis of the embryo proper lengths for the indicated genotypes (N≥20) at the globular stage (Welch’s ANOVA followed by Tukey’s test, statistically significant differences compared to WT are shown, *** is significant at p<0.001, ** is significant at p<0.01, * is significant at p<0.05). Results from both statistical analyses (B,C) are in Tables S3 and S4, respectively. In box plots the middle line in the box represents median, the × shows mean, the bottom line depicts the 1st quartile, while the top line describes the 3rd quartile; the vertical lines (whiskers) extend to minimum and maximum value within the 1.5× interquartile range (distance between the 1st and the 3rd quartile); points outside of the whiskers mark outliers (values outside of the 1.5× interquartile range).

Shorter embryo proper length was found in *yda-1* and *ΔNyda* mutants (Fig.1C), in contrast to *hsp90.2* and *hsp90^RNAi^* that showed longer embryo proper, whereas in *hsp90.1* there was no significant change (Fig.1C). The length of embryo proper was restored to wild-type-like values in *hsp90^RNAi^ yda-1 and hsp90^RNAi^ ΔNyda* double mutants, while *hsp90.1 yda-1* and *hsp90.2 yda-1* displayed longer embryo propers (Fig.1C). In *hsp90.2 ΔNyda*, we also observed increased embryo proper length compared to single *yda-1* and *ΔNyda* and the wild-type (Fig.1C). Although the embryo proper was not significantly longer in *hsp90.1 ΔNyda* compared to the wild-type, it was significantly longer than in the *ΔNyda* single mutant (Fig.1C). These results show that *HSP90.1* depletion or reduced activity of HSP90s suppresses the suspensor and embryo proper phenotype in both *yda-1* and *ΔNyda* mutants, while HSP90s are involved in the control of both basal and apical embryo development. This suggests an epistatic relationship between *HSP90.1* and *YDA*.

To distinguish the type of the epistatic effect of *HSP90s* on *YDA* regarding embryo suspensor length, we calculated the epistatic values of double mutants. The epistatic effect on a specific trait can be synergistic, when the phenotypic value of double mutants is higher than the linear combination of single mutants. When the phenotypic value of double mutants is lower than the linear combination of single mutants it is described as antagonistic epistasis (Hartman *et al*., 2001; Segrè *et al*., 2005). The *hsp90 yda-1* double mutants display positive epistasis values that correspond to synergistic epistasis, while *hsp90 ΔNyda* double mutants show negative epistasis values corresponding to antagonistic epistasis (Supplemental Fig.S2). Epistatic effects can shift from synergistic to antagonistic depending on the activity of the protein involved, like in the case of gain- and loss-of-function mutations or the tissue where this activity is measured (Li *et al*. 2018). The epistatic effect of *hsp90.1, hsp90.2* and *hsp90^RNAi^* towards *yda-1* regarding embryo suspensor elongation was different, while the epistatic effect of *hsp90.1* and *hsp90.2* to the constitutively active form of YDA in *ΔNyda* mutants was similar (Supplemental Fig.S2). Taking together, our analysis demonstrates an epistatic interaction between HSP90s and YDA in the regulation of suspensor length in developing embryos.

### Genetic impairment of HSP90s and/or YDA leads to aberrant CDPs during early embryo development

Embryo development relies on a complex transcriptional network that regulates asymmetric cell divisions and embryo patterning. Since early stages of embryo development are characterized by an amazing precision of cell divisions (Yoshida *et al*., 2014), to investigate the role of HSP90 and YDA interplay in the orientation of CDPs, we analyzed cell division patterns in early embryonic stages of *hsp90s* and *yda* single and double mutants. Mis-patterned CDPs were observed in both the proembryo and suspensor of *hsp90.1, hsp90.2, hsp90^RNAi^* leading to abnormal cell division patterns. The *hsp90.1* mutant showed the highest frequency of these disorders (Fig.2). Consistent with previously published results (Lukowitz *et al*., 2004) *yda-1* mutants displayed asymmetric and ectopic cell divisions in both basal and apical parts, while *ΔNyda* embryos displayed asymmetric divisions but no ectopic ones (Fig.2). Irregular cell divisions were detected in *mpk3-1, mpk6-2* (for roots see also Müller *et al*., 2010) and *mpk6AEF* (a non-phosphorylatable form of *MPK6;* Bush and Krysan, 2007) embryos, leading, however, to less severe phenotypes than the *yda-1* mutants (Supplemental Fig.S3).

**Fig.2.**
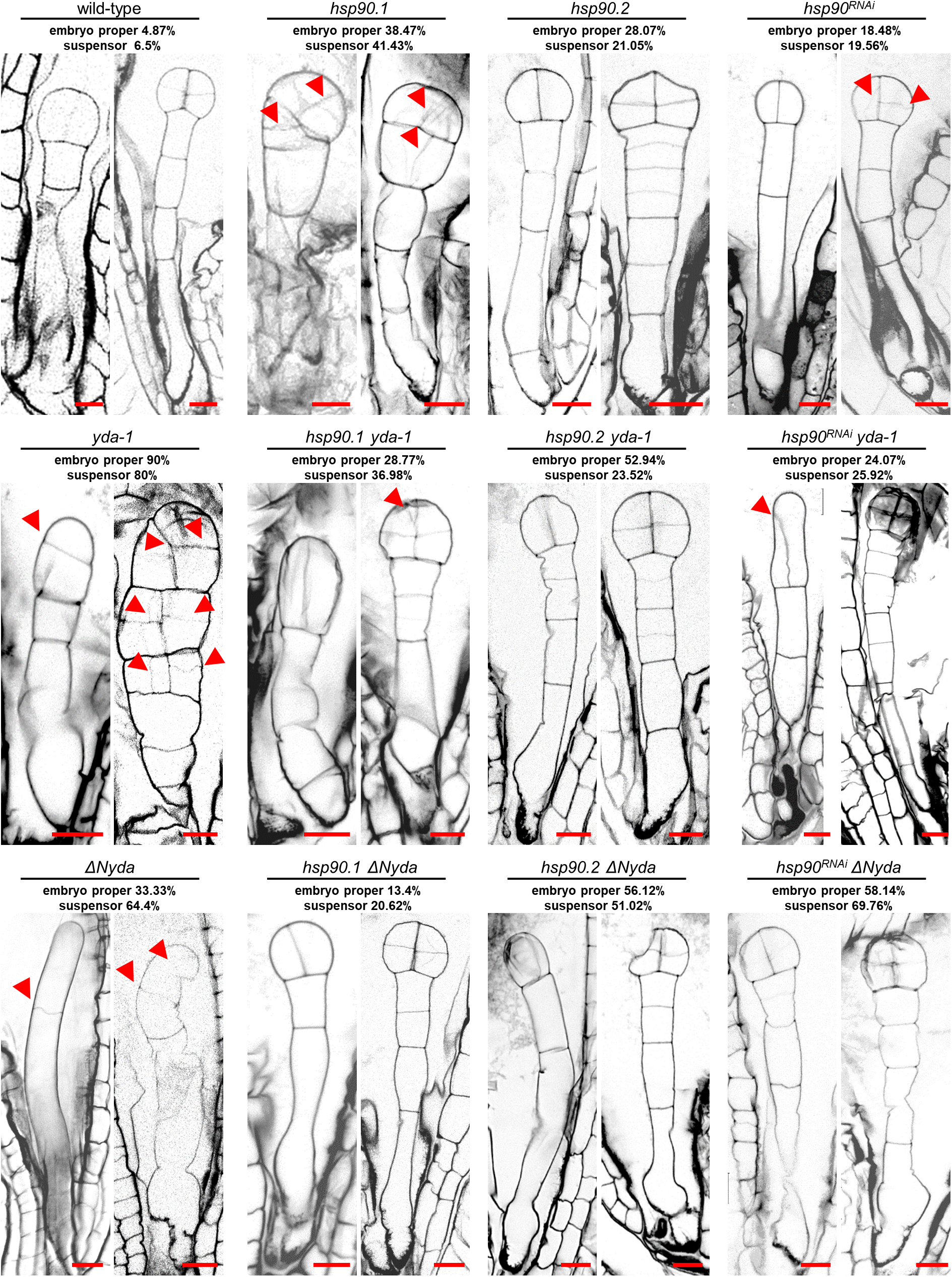
Genetic depletion of HSP90s restores defects in cell division plane orientation in the early embryogenesis of *yda* mutants. Representative images of early embryonic development at the 2-to 4-cell stage and 8-cell stage for each of the indicated phenotypes. Developing embryos were visualized with propidium iodide staining (shown in inverted black-white mode). Red arrowheads point to abnormally positioned cell walls. The frequencies of the observed disorders in division patterns in embryo suspensor and abnormalities in embryo proper of the different genotypes are presented above the corresponding images (N≥20). Scale bars, 10 μm.

Notably, HSP90s genetic depletion suppressed the CDP defects in *yda-1* embryos, as we detected fewer ectopic cell divisions in *hsp90.1 yda-1* double mutants (Fig.2). In *hsp90.2 yda-1* we observed atypical CDPs mainly in the apical cell lineage, while lower frequency of impaired CDPs was observed both in the basal and apical cell lineage in *hsp90^RNAi^ yda-1* (Fig.2). Similar trend was observed in *hsp90 ΔNyda* double mutants, where the genetic depletion of HSP90s resulted in fewer CDP defects. This partial rescue was especially prominent in the *hsp90.1 ΔNyda* double mutant (Fig.2). Comparing the frequency of abnormal CDPs in the wild-type on one side and the *yda-1* single mutant on the other side, the percentages of irregular cell division in *hsp90* and *yda-1* double mutant suspensors and embryo propers resembled the frequencies of defective cell divisions in *hsp90* single mutants (Fig.2). These results suggest that HSP90s regulate YDA signaling cascade during the establishment of both apical and basal embryonic cell fates, while their depletion can restore fully or partially the severe defects in CDP orientations during early embryo development in *yda* mutants.

### Mispatterned *WOX8* expression in *hsp90, yda-1* and *ΔNyda* mutants

*WOX* genes are specifically expressed in developing embryos but not in other tissues within siliques. *WOX8* and its paralogue *WOX9* act redundantly in suspensor and proembryo development (Breuninger *et al*., 2008; Ueda *et al*., 2011), and are both required for *WOX2* expression. Since the YDA cascade controls the expression of *WOX8* (Ueda *et al*., 2017), we analyzed the expression of *WOX8* in *hsp90, yda-1, ΔNyda, mpk3-1, mpk6-2* and *mpk6AEF* by using *proWOX8:NLS-YFP* construct. By analysing the percentage of developing embryos expressing *proWOX8:NLS-YFP* molecular marker during early and later stages of embryo development we showed that in wild-type plants *WOX8* is expressed almost stably, since the early stages of development till the dermatogen stage (Supplemental Fig.S4A). All *hsp90* mutants showed lower percentage of developing embryos expressing *proWOX8:NLS-YFP* in the early stages of embryo development. In *hsp90.1* and *hsp90^RNAi^* it remained low in later stages of embryo development; however, in *hsp90.2* it approached the wild-type levels in later stages (Supplemental Fig.S4A). Analysis of *proWOX8:NLS-YFP* expression in *mpk3-1* showed that 78% of early staged embryos expressed the marker, while this percentage increased to 86% later in development (Supplemental Fig.S4A). In *mpk6-2* and *mpk6AEF* the expression percentage was lower than in *mpk3-1* in early stages. The expression in later stages was similar to wild-type, especially in the *mpk6-2* (Supplemental Fig.S4A). A similar trend was observed in *yda-1* mutant. In contrast, the expression of *proWOX8:NLS-YFP* in *ΔNyda* was close to wild-type levels throughout embryo development (Supplemental Fig.S4A).

We showed very low expression of *WOX8* marker in the suspensor cells of young embryos in *hsp90.1* and enhancement later in development; however, the absence of *WOX8* expression in the suspensor cells close to the hypophysis was typical in *hsp90.1* mutant embryos (Fig.3 and Supplemental Fig.S5). In *hsp90.2* embryos, the expression of *WOX8* was higher than in *hsp90.1*, but the aberrant expression patterns were similar (Fig.3 and Supplemental Fig.S5). In *hsp90^RNAi^* the abnormal expression of *proWOX8:NLS-YFP* was induced, as in most of the embryos *WOX8* expression was confined at the basal cells of the suspensor (Fig.3 and Supplemental Fig.S5). These findings suggest that HSP90s are involved in the spatiotemporal regulation of *WOX8* expression during early embryogenesis.

**Fig.3.**
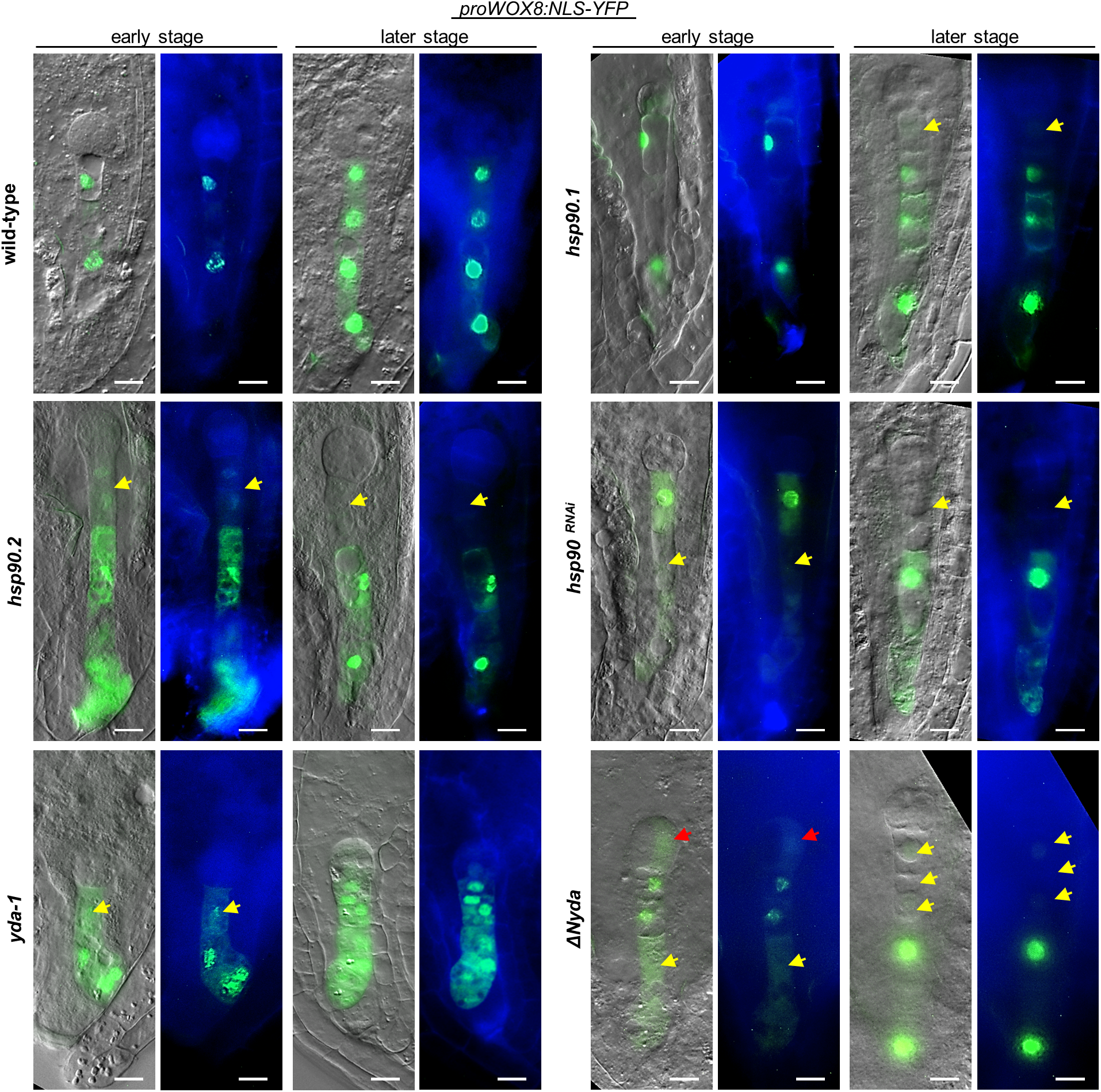
Impaired function of HSP90s and YDA leads to aberrant transcriptional activation of *WOX8* affecting basal cell fate determination. Analysis of *proWOX8:NLS-YFP* localization and signal distribution during early stage and later stage of early embryogenesis in the indicated genotypes. From left to right: merged images of DIC with fluorescent YFP signals and SCRI Renaissance 2200 staining with fluorescent YFP signals. Yellow arrows point to missing expression of *proWOX8:NLS-YFP* in the suspensor cells, red arrows indicate ectopic expression in embryo proper. Scale bars, 10 μm.

Further analysis showed induced *WOX8* expression in the suspensor of *mpk3-1* embryos, although in the early developmental stages basal cells of the suspensor express less the marker (Supplemental Fig.S6A, S7). The expression of *WOX8* was hardly visible in the suspensors of *mpk6-2* embryos throughout the development, but in *mpk6AEF* where the activation of MPK6 through phosphorylation is blocked, the ectopic expression of *WOX8* in the apical lineage was quite profound (Supplemental Fig.S6A, S7). In *yda-1* embryos, *proWOX8:NLS-YFP* was overexpressed in all the developmental stages (Fig.3 and Supplemental Fig.S7). In contrast, *ΔNyda* mutant showed hardly detectable expression of *WOX8* in the suspensor and ectopic expression in the apical lineage during the early stages of embryo development (Fig.3 and Supplemental Fig.S7). *WOX8* levels increased later in development, but they were significantly lower compared to the wild-type, while most of suspensor cells close to the hypophysis did not express *WOX8* in *ΔNyda* embryos (Fig.3). Quantitative analysis of *WOX8* expression in *hsp90, yda-1, ΔNyda, mpk3-1, mpk6-2* and *mpk6AEF* mutants showed increased frequencies of embryos displaying missing signal of *proWOX8:NLS-YFP* expression in all the cells of the suspensor or ectopic signal in the apical part (Supplemental Fig.S6B and C). The analysis of *WOX8* expression in *mpk3-1* and *mpk6-2* suggests different roles of MPK3 and MPK6 in the transcriptional activation of *WOX8* indicating that activation of MPK6 through phosphorylation restricts *WOX8* expression in the basal cell lineage. Moreover, YDA depletion or overactivation have opposite effects on the control of *WOX8* expression. Thus, YDA signaling cascade is involved in the spatiotemporal transcriptional activation of *WOX8*.

### Impaired HSP90s function downregulates *WOX2* expression

To test the impact of *WOX8* transcriptional misregulation on *WOX2* expression, we analysed *proWOX2:NLS-YFP* expression in the respective genetic backgrounds. In wild-type plants there was a delay in the expression of *proWOX2:NLS-YFP* marker, as we could detect fluorescent signals in 84% of early embryos (Supplemental Fig.S4B), which was increased later in development. In all *hsp90* mutants, we observed lower frequencies of *proWOX2:NLS-YFP* expressing embryos during the early stages which were increased in later developmental stages, but still remained lower compared to the wild-type (Supplemental Fig.S4B). The expression frequency of *proWOX2:NSL-YFP* was very low in early stages of *mpk3-1* embryos (38%) and significantly increased later in development (Supplemental Fig.S4B). The percentage of *mpk6-2* and *ΔNyda* embryos displaying fluorescent signal of *WOX2* marker was stable during embryo development, however, it was lower than in the wild-type. Finally, the percentage of embryos expressing the *proWOX2:NLS-YFP* marker in *mpk6AEF* followed a similar trend with *hsp90* mutants (Supplemental Fig.S4B).

Comparison of the percentages of embryos expressing *proWOX8:NLS-YFP* versus *proWOX2:NLS-YFP* showed a positive correlation in the expression of the two markers in wild-type corroborating published data (Breuninger *et al*., 2008). *WOX2* expression in later developmental stages in *hsp90.1, hsp90^RNAi^* and *mpk6-2* mutants did not follow the expression of *WOX8* (Supplemental Fig.S4), suggesting additional roles for HSP90.1 and MPK6 in the regulation of *WOX2* expression.

Next, we analysed *proWOX2:NLS-YFP* expression pattern in *yda-1* and *ΔNyda* that exhibit opposite traits concerning the apical-basal axis formation. Semiquantitative analysis of signal intensity showed that *WOX2* expression was established in the apical cell lineage during early embryogenesis (2-cell embryos) and the signal intensity was similar in later developmental stages in wild-type (Fig.4A). In *yda-1* embryos, we observed increased expression of *proWOX2:NLS-YFP* in early developmental stages and strong ectopic expression in suspensor cells from which eventually the hypophysis will develop (Fig.4B). Such ectopic expression of *WOX2* was accompanied by irregular cell divisions (Fig.4B). In *ΔNyda* embryos the expression of *proWOX2:NLS-YFP* was very low, while ectopic expression of the marker was detected in suspensor cells undergoing irregular and/or shifted cell divisions (Fig.4C). Our results suggest that YDA controls the levels and the spatiotemporal expression of *WOX2*, since depletion or constitutive activation of YDA have opposite effects on *WOX2* transcriptional activation. *hsp90^RNAi^* showed *WOX2* transcriptional activation in the suspensor cells (Fig.4D). Similarly, *hsp90.1* and *hsp90.2* embryos revealed enhanced and ectopic expression of *WOX2* in the suspensor (Supplemental Fig.S8A) suggesting that HSP90s are essential for cell fate determination. Finally, ectopic expression of *proWOX2:NLS-YFP* was also detected in the suspensors of *mpk3-1* embryos (Supplemental Fig.S8B).

**Fig.4.**
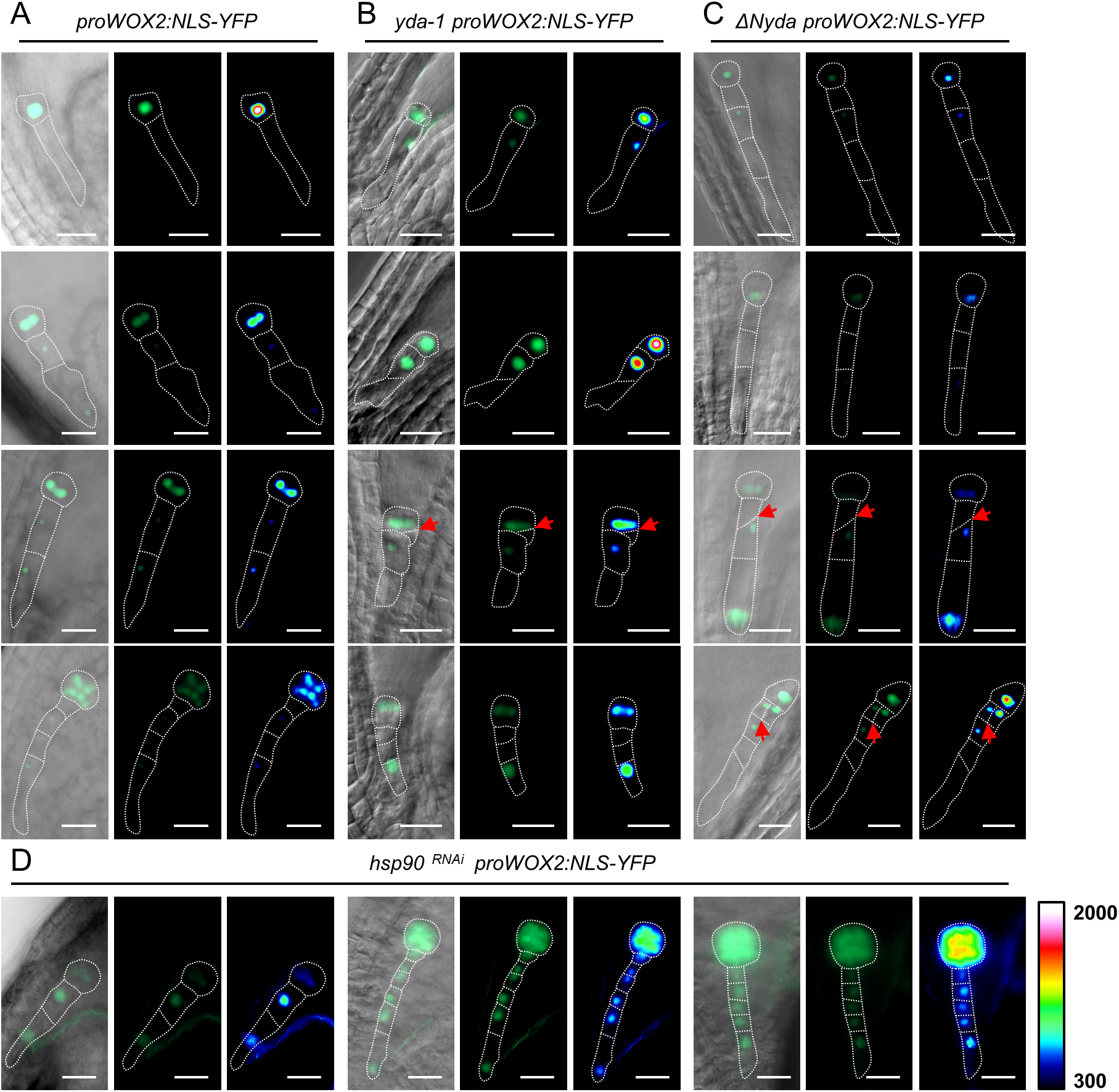
Aberrant function of HSP90s and YDA impacts *WOX2* expression in early embryogenesis. (A-D) Expression pattern of *proWOX2:NLS-YFP* marker in developing embryos of wild-type (A), *yda-1* (B), *ΔNyda* (C) and *hsp90^RNAi^* (D) mutants. From left to right: merged images of DIC with fluorescent YFP signals, fluorescent YFP signals, and semi-quantitative fluorescence intensity evaluation of *WOX8* expression levels using pseudo colour-coded range from the scale, where dark blue represents zero intensity (300 in arbitrary units) and white represents maximum intensity (2000 in arbitrary units). Red arrows point to abnormally positioned cell walls. Scale bars, 20 μm.

### Impaired function of HSP90s or YDA signaling cascade leads to altered auxin maxima

Since WOX transcriptional machinery has been implicated in the establishment of localized auxin response during embryo development (Breuninger *et al*., 2008), we analyzed auxin maxima using *proDR5:GFP* construct in developing embryos of *hsp90, yda-1, ΔNyda, mpk3-1, mpk6-2* and *mpk6AEF* mutants. Analysis of heart and late heart embryonic stages of *hsp90.1, hsp90.2* and *hsp90^RNAi^* showed deregulations of *proDR5:GFP* expression in the root and cotyledon embryo poles and decreased auxin maxima suggesting alterations in the auxin distribution (Fig.5A). The greatest reduction in *proDR5:GFP* signal was observed in *hsp90^RNAi^*, as demonstrated by the semiquantative evaluation of the fluorescent *proDR5:GFP* in developing embryos (Fig.5A). Quantification of basal to apical *proDR5:GFP* signal intensity ratio and analysis of *proDR5:GFP* expression patterns provided further evidence for disturbed auxin distribution in the embryos of *hsp90* mutants (Fig.5B and C).

**Fig.5.**
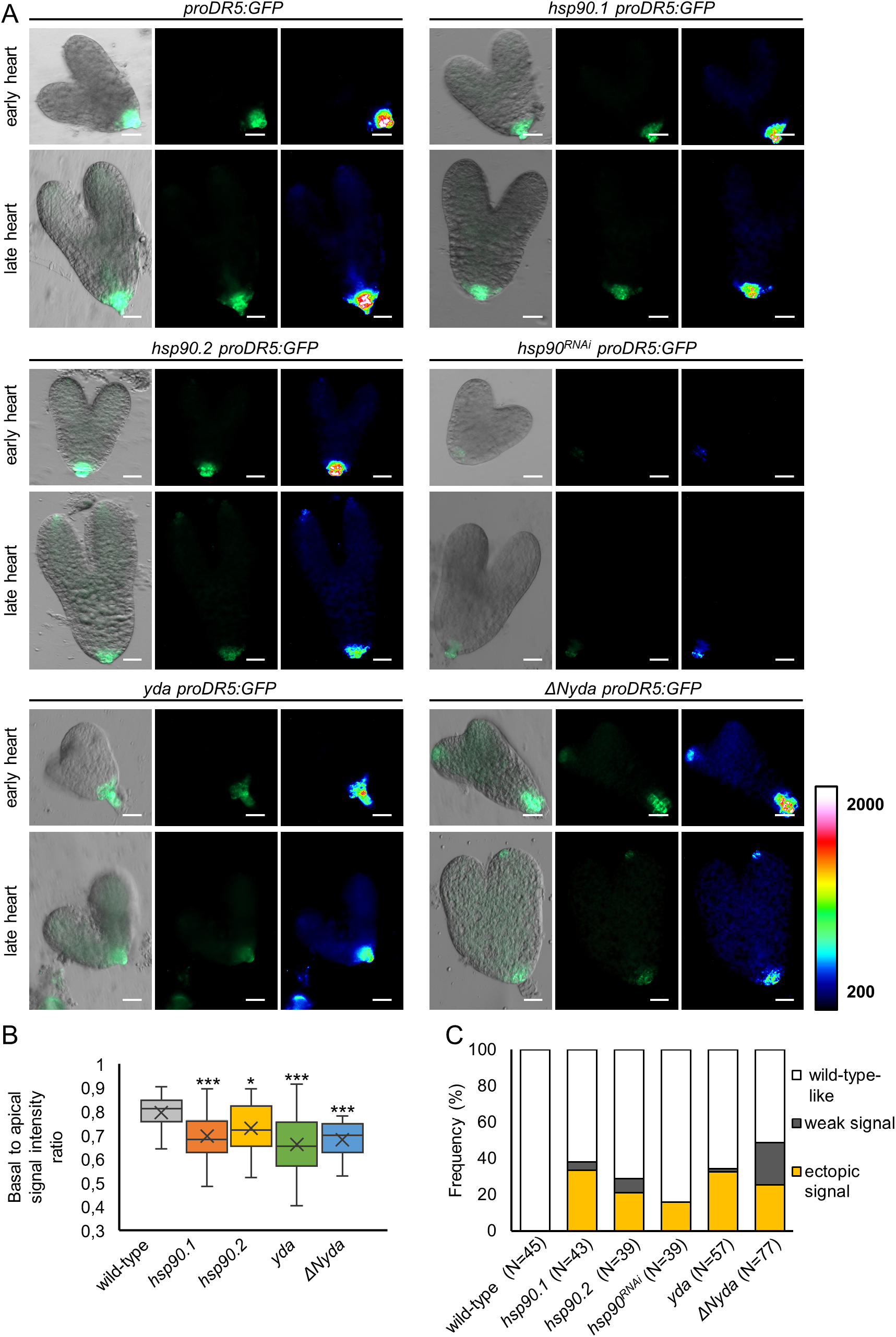
HSP90s and YDA affect auxin maxima patterns during embryo development. Auxin maxima were visualized in lines expressing *proDR5:GFP*. (A) Microscopic analysis of *proDR5:GFP* signal distribution in late embryonic development (early and late heart stages) is shown for the indicated genotypes. From left to right: merged images of DIC with fluorescent GFP signals, fluorescent GFP signals, and semi-quantitative fluorescence intensity evaluation of *proDR5:GFP* expression levels using pseudo color-coded range from the scale, where dark blue represents zero intensity (200 in arbitrary units) and white represents maximum intensity (2000 in arbitrary units). Scale bars, 20 μm. (B) The ratio of basal to apical fluorescence signal intensity showing auxin distribution within embryos (N>27; Kruskal-Wallis test, statistically significant differences compared to WT are shown, * is significant at p<0.05, *** is significant at p<0.001). The results of statistical analysis are in Table S5. (C) Quantitative analysis of defects in auxin maxima localization in the indicated genotypes (N=number of observations).

In *mpk3-1* and *mpk6-2* we observed accumulation of active auxin both in the cotyledon and root poles of embryos (Supplemental Fig.S9A), while abnormal auxin distribution between the basal and apical axis of the embryo was found in *mpk6-2* embryos (Supplemental Fig.S9B). The opposite effects of *mpk6-2* and *mpk6AEF* on *proDR5:GFP* intensity suggest that the establishment of localized auxin response likely depends on the activation of MPK6 (Supplemental Fig.S9A). In *yda-1* embryos there were increased auxin maxima in the root pole both in early and late heart stages, but no detectable signal in the cotyledon poles, while aberrant expression of the marker was also detected (Fig.5A and C). In *ΔNyda*, weak auxin accumulation was observed in the root pole of embryos, whereas some aberrant signal was found in the cotyledon poles (Fig.5A and C). These alterations were further supported by the quantification of basal to apical fluorescent signal intensity ratio of *proDR5:GFP* in *yda-1* and *ΔNyda* embryos (Fig.5B). Interestingly, *HSP90.1* depletion in *yda-1* mutant compensated auxin maxima in both cotyledon and root poles of the *hsp90.1 yda-1* embryos, however, the auxin maxima were lower compared to wild-type (Supplemental Fig.S10). Taken together, our data show that HSP90s and YDA are important for the establishment of localized auxin maxima during embryonic development.

### HSP90s and MPKs influence histone H3 deposition at WRKY binding sites in the promoters of *WOX* genes

Disturbed expression of *WOX8* and *WOX2* did not always correlate with YDA activity, since constitutively active YDA cascade in *ΔNyda*, which enhances WRKY2 activation through phosphorylation (Ueda *et al*., 2017), did not result in higher levels of *WOX8* transcripts. These findings imply the involvement of additional regulatory mechanisms contributing to the transcriptional regulation of *WOX* genes.

The critical role of chromatin dynamics and modifications in the regulation of transcription during embryogenesis was reported previously (Shindo and Amodeo, 2019; Bouyer *et al*., 2011). Since genetic depletions of HSP90s led to the rescue of severe phenotypic defects in *yda-1* and *ΔNyda* embryos, we investigated whether HSP90s participate in the epigenetic control of *WOX8, WOX9* and *WOX2*. Changes in the composition of chromatin could contribute to alterations in chromatin structure that permit the association of transcription factors at the corresponding binding sites in the promoters of their target genes. To understand the mechanisms controlling the spatiotemporal patterns of expression of *WOX* genes, we investigated chromatin dynamics by testing histone H3 occupancy at WRKY binding sites residing in the promoters of *WOX* genes by performing H3-ChIP qPCR analysis in *hsp90.1, ΔNyda* and *hsp90.1 ΔNyda* mutants, as WRKY2 is one of the major transcription factors regulating *WOX8* expression (Ueda *et al*., 2017). To this end, we performed H3-ChIP qPCR analysis at WRKY binding sites residing upstream the transcription initiation site of the target genes in wild-type, *hsp90.1, ΔNyda* single mutants and *hsp90.1 ΔNyda* double mutant (Fig.6A). *LEAFY COTYLEDON 2 (LEC2*) promoter was used as a positive control for ChIP, while *FUSCA 3 (FUS3*) promoter, recently reported to be regulated by WRKY transcription factors during seed maturation, was used as an additional positive control (Roscoe *et al*., 2018). H3 ChIP qPCR analysis showed that the selected genomic regions, namely WRKY boxes (Fig.6A), correspond to nucleosomal sites as it was evident by the binding of H3 to these loci in wild-type plants (Supplemental Fig.S11A). Then, we tested H3 occupancy at the same genomic sites among different genetic backgrounds (Fig.6B). Our results showed that HSP90.1 depletion and constitutive activation of YDA in *ΔNyda* increased H3 occupancy at WRKY binding sites at almost all tested promoters, while in *hsp90.1 ΔNyda* double mutant the abundance of H3 was compensated to almost wild-type levels in the promoters of *WOX8, WOX9* and *WOX2* genes (Fig.6B). Histone H3 distribution at *FUS3* promoter in *hsp90.1 ΔNyda* double mutant did not follow the trend of *WOX* genes (Fig.6B). These results strongly suggest that impaired function of HSP90.1 and constitutive activation of YDA increase the H3 pools and, thus, nucleosome occupancy at the WRKY binding sites of *WOX* gene family promoters.

**Fig.6.**
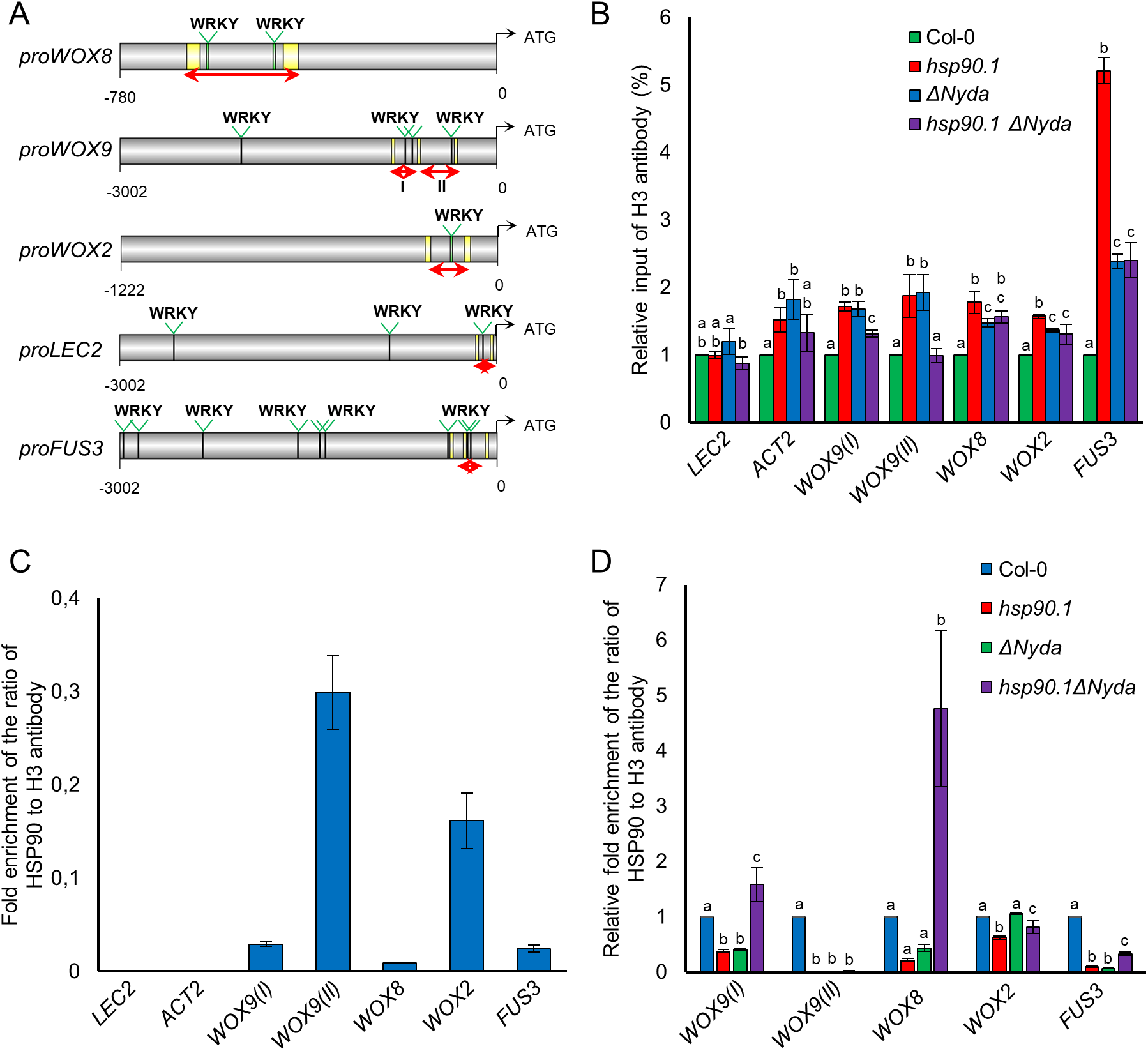
HSP90s are present at the WRKY binding sites regulating H3 occupancy in the promoters of genes involved in body axis formation during early embryogenesis. (A) Schematic depiction of WRKY binding sites (TTGACT/C) at the promoters of denoted genes. For the analysis, AtcisDB – Arabidopsis cis-regulatory element database (https://agris-knowledgebase.org/AtcisDB/) was used. Red arrows at the promoters show the amplified genomic regions. (B) Relative H3 input (%) at WRKY boxes at *LEC2*, *ACT2, WOX9*, *WOX8*, *WOX2* and *FUS3* promoters of the indicated genotypes. Data are presented as means±SD (N≥three biological and technical triplicates; ANOVA followed by Scheffe’s test was applied for statistical evaluation; letters in the graph are shared by groups without statistically significant differences at the 0.05 probability level; results of statistical analysis are in Table S6). (C) ChIP using HSP90 antibody followed by qPCR analysis of WRKY binding sites at the promoters of *LEC2*, *ACT2, WOX9*, *WOX8*, *WOX2* and *FUS3* in wild-type. (D) Relative fold enrichment of HSP90s normalized to H3 in *hsp90.1, ΔNyda* and *hsp90.1 ΔNyda*. Data are presented as means±SD (N≥three biological and technical triplicates; ANOVA followed by Scheffe’s test was applied for statistical evaluation; letters in the graph are shared by groups without statistically significant differences at the 0.05 probability level; results of statistical analysis are in Table S7).

### HSP90s are involved in the epigenetic control of *WOX* family genes

To test the presence of HSP90s at the specific genomic loci, we performed HSP90-ChIP qPCR for the same genomic loci as in the abovementioned experiment (Fig.6A). Our results showed a higher abundance of HSP90s (normalized to H3 levels) at the WRKY binding sites in the promoters of *WOX* family genes and *FUS3* in comparison to the positive control *LEC2* in the wild-type (Fig.6C). We found significantly decreased levels of HSP90 at the WRKY binding sites in the promoters of *WOX8, WOX9, WOX2* and *FUS3* in *hsp90.1* (Fig.6D), while in the promoter of the positive control, *LEC2*, HSP90 levels were increased (Supplemental Fig.S11B). A similar trend was observed in the promoters of *WOX8, WOX9* and *FUS3* genes in *ΔNyda* mutants except of *WOX2* promoter where the levels of HSP90s at the WRKY binding sites were similar to the wild-type (Fig.6D). In the *hsp90.1 ΔNyda* double mutant, there was an increase in HSP90s levels compared to single mutants (Fig.6D). Thus, these findings suggest a negative correlation between constitutive activation of YDA and HSP90 levels at the WRKY binding sites of the studied genes.

## Discussion

Developmental and environmental cues integrate with essential signaling components during embryogenesis to secure the successful development of the offspring. Formation of the main body axis during embryo development requires spatiotemporal regulation of gene expression in embryonic and suspensor cells, which acquire distinct properties essential for the establishment of different cell fates. Our results suggest that the genetic interaction of *HSP90s* with *YDA* affects asymmetric cell division patterns and promotes directional growth in the early developing embryo. This is associated with the modulation of the spatiotemporal restriction of *WOX8* and *WOX2* expression, which is mediated possibly through the epigenetic regulation of the transcriptional networks marking the onset of embryo development.

Genetic depletion of HSP90 family members restored the severe embryo phenotypes of both loss-of-function *yda-1* mutants and gain-of-function *ΔNyda* mutants, which was evident by the recovery of suspensor and embryo proper elongation to wild-type levels and the restoration of cell division defects in *hsp90 yda-1* and *hsp90 ΔNyda* double mutants. These genetic studies suggest that HSP90s act epistatically to YDA.

A combination of two principle mechanisms, such as segregation of intrinsic determinants during division and extrinsic stimuli define cell division patterns during embryo development (ten Hove *et al*., 2015). The homeodomain transcription factors WOX2 and WOX8 play crucial roles as cell-lineage-specific developmental regulators in the establishment of apical-basal axis formation (Breuninger *et al*., 2008; Ueda *et al*., 2011). The independent expression of *WOX2* and *WOX8* after the asymmetric division of the zygote establishes different transcriptional landscapes in the resulting daughter cells. These different transcriptional programs also affect auxin distribution in the developing embryo (Breuninger *et al*., 2008; Ueda *et al*., 2011). Genetic depletion of WOX8/9 suppresses the expression of *WOX2* required for shoot apex patterning (Breuninger *et al*., 2008). YDA pathway, acting as an extrinsic receptor-mediated cue, is involved in the transcriptional control of *WOX8* by integrating parental signals to activate WRKY2 transcriptional function (Ueda *et al*., 2017). In this study, we report that aberrant expression of *WOX8* in *hsp90, yda, mpk3 and mpk6* mutants are directly related to the ectopic expression of *WOX2* in the basal cell lineage and to the disturbed distribution of auxin during embryonic development.

Most formation and transition steps in the early embryo of Arabidopsis depend on auxin biosynthesis, transport and signaling (Möller and Weijers, 2009). HSP90s interact with the N-terminal regulatory domain of YDA and affect the activation through phosphorylation of downstream targets, like MPK3 and MPK6 (Samakovli *et al*., 2020b). MPK6 was found to phosphorylate PIN-FORMED 1 (PIN1) auxin efflux transporter regulating polar auxin transport affecting a series of auxin-mediated developmental processes (Jia *et al*., 2016). Ectopic cell divisions (Müller *et al*., 2010) and higher auxin levels were observed in both *yda-1* and *ΔNyda* mutants attributed to up-regulated auxin biosynthesis (Smékalová *et al*., 2014). Also, WOX8 and WOX9 control the activation of WOX2 to mediate auxin patterning through the regulation of PIN1 (Haecker *et al*., 2004; Wu *et al*., 2007; Breuninger *et al*., 2008). Our results show that impaired function of HSP90s results in lower auxin maxima suggesting that the molecular chaperones are involved not only in the auxin signaling through their interaction with TIR1 (TRANSPORT INHIBITOR RESPONSE 1) auxin receptor (Wang *et al*., 2016; Watanabe *et al*., 2016), but also in the polar distribution of auxin. Problems in the establishment of localized auxin maxima response were encountered in both *mpk3-1* and *mpk6-2* mutants, which displayed increased localized auxin maxima during embryogenesis. By contrast, the auxin maxima were significantly reduced in *mpk6AEF*. Aberrant auxin distribution was also observed in *yda-1* and *ΔNyda* embryos.

Epigenetic control is essential for the regulation of gene transcription during plant development (Köhler *et al*., 2012). Polycomb group proteins (PcG) proteins regulate gene expression by epigenetically modifying chromatin (Schwartz and Pirrota, 2013). The core PRC2 complex is conserved between animals and plants (Xiao and Wagner, 2015). Many studies point out that PRC2 plays a critical role during seed development in regulating the cascade of transcription factors and other genes involved in tissue specification (Mozgova *et al*., 2015; Xiao and Wagner, 2015). Transcription factors from the WOX family have been identified as targets of PRC2 (Bouyer *et al*., 2011). Besides, *FUS3*-related transcriptional control is also mediated by PRC2 through the deposition of H3K27me3 (Roscoe *et al*., 2018).

Analysis of *WOX8* and *WOX2* spatiotemporal expression patterns in developing embryos of *hsp90, yda, mpk3 and mpk6* mutants showed aberrant expression for both genes, suggesting that HSP90s and YDA signaling cascade are essential for basal and apical cell fate determination. The molecular link between YDA cascade and the regulation of *WOX8* expression is WRKY2 transcription factor, which is phosphorylated by MPK3 and MPK6 to mediate the transcriptional activation of *WOX8* (Ueda *et al*., 2017). A recent study in *Manihot esculenta* showed that HSP90s can promote directly the transcriptional activation of *NINE-CIS-EPOXYCAROTENOID DIOXYGENASE 5* (*MeWRKY20)* through WRKY-binding sites residing in the promoter controlling the expression of this gene, which encodes a key enzyme of ABA biosynthesis, to regulate drought stress response (Wei *et al*., 2019). Our analysis showed that the genetic interplay between HSP90s and YDA influences H3 pools and nucleosome occupancy at the WRKY binding sites in the promoters of *WOX8, WOX9* and *WOX2* genes. Additionally, we showed that HSP90 are present in the WRKY boxes of *WOX* genes implicating the transcriptional activation through WRKY transcription factors regulating the spatiotemporal restriction of *WOX8* and *WOX2* expression.

HSP90s act as components of chaperone complexes that facilitate their binding to specific client proteins. The broad range of HSP90 interactors places these molecular chaperones at the position of a central hub in many signaling pathways in Arabidopsis (Tichá *et al*., 2020). Recent studies also highlighted the capacity of HSP90s to form molecular scaffolds and maintain gene circuitries during flowering (Margaritopoulou *et al*., 2016). Moreover, involvement of HSP90s in auxin and brassinosteroid signaling pathways (Samakovli *et al*., 2014; 2020a; Shigeta *et al*., 2015; Wang *et al*., 2016; Watanabe *et al*., 2016) suggests a critical role in the control of plant development. Indeed, the expression of cytosolic HSP90 members is strictly regulated during embryo development (Prasinos *et al*., 2005), and *hsp90* mutants display embryo abortion phenotypes (Samakovli *et al*., 2007), and chloroplast biogenesis-associated HSP90.5 is essential for embryogenesis (Feng *et al*., 2014).

The regulation of transcriptional machinery by HSP90s takes place at different levels such as the activation/inactivation of certain transcription factors, the modulation of the activity of chromatin remodelling factors, like DNA methyltransferases and histone deacetylases and the removal of histones from the promoters of specific genes (Khurana and Bhattacharyya, 2015). HSP90s participate in stomatal ontogenesis by regulating YDA localization and polarity, the activation of downstream signaling components, and the transcriptional activity of SPCH, the master transcription factor controlling stomatal development (Samakovli *et al*., 2020b), whereas they regulate stomatal differentiation by controlling the levels of SPCH and MUTE transcription factors through the activation of the YDA signaling cascade (Samakovli *et al*., 2020c).

HSP90s are involved in the epigenetic regulation of gene expression, since they participate in chromatin accessibility through interactions with chromatin remodelling complexes that activate or repress gene expression, including SMYD3 (SET and MYND domain-containing protein 3), trithorax/Mixed Lineage Leukemia (MLL) proteins, RSC (Remodelling the Structure of Chromatin) complex and PRC2 complex catalytic subunit EZH2 (Enhancer of Zeste Homolog 2) in humans (Fiskus *et al*., 2009; Tariq *et al*., 2009; Brown *et al*., 2015; Echtenkamp *et al*., 2016). HSP90s bind close to the transcription initiation site of one-third of protein-coding genes in ChIP experiments performed in Drosophila (Yoveva and Sawarkar, 2018). Taken together, these reports highlight emerging roles of HSP90s in the epigenetic regulation of gene expression.

Here, we demonstrate the dynamic role of HSP90s in the spatiotemporal restriction of *WOX8* and *WOX2* expression during early embryonic development of Arabidopsis as described in the proposed model (Fig.7). HSP90s and YDA cascade regulate nucleosome occupancy on the WRKY binding sites in the promoters of *WOX8*, *WOX9* and *WOX2* genes, as disturbed function of HSP90s or overactivation of YDA results in increased deposition of nucleosomes in the respective genomic loci. Nucleosome occupancy at the promoters of individual genes is inversely proportional to their transcription rate as a possible result of transcription factors displacing nucleosomes (Ercan *et al*., 2004). Constitutive activation of YDA leads to reduced HSP90 levels on the WRKY boxes of *WOX* genes. Therefore, an HSP90-mediated nucleosome depletion from WRKY regulatory elements could be suggested as a new mechanism controlling the spatiotemporal restriction of *WOX8* and *WOX2* expression during early embryogenesis (Fig.7). Finaly, our results show a new role for the crosstalk between HSP90s and YDA signaling cascade in the control of the transcriptional networks that determine apical and basal cell fates shaping the formation of main body axis during early embryo development in Arabidopsis.

**Fig.7.**
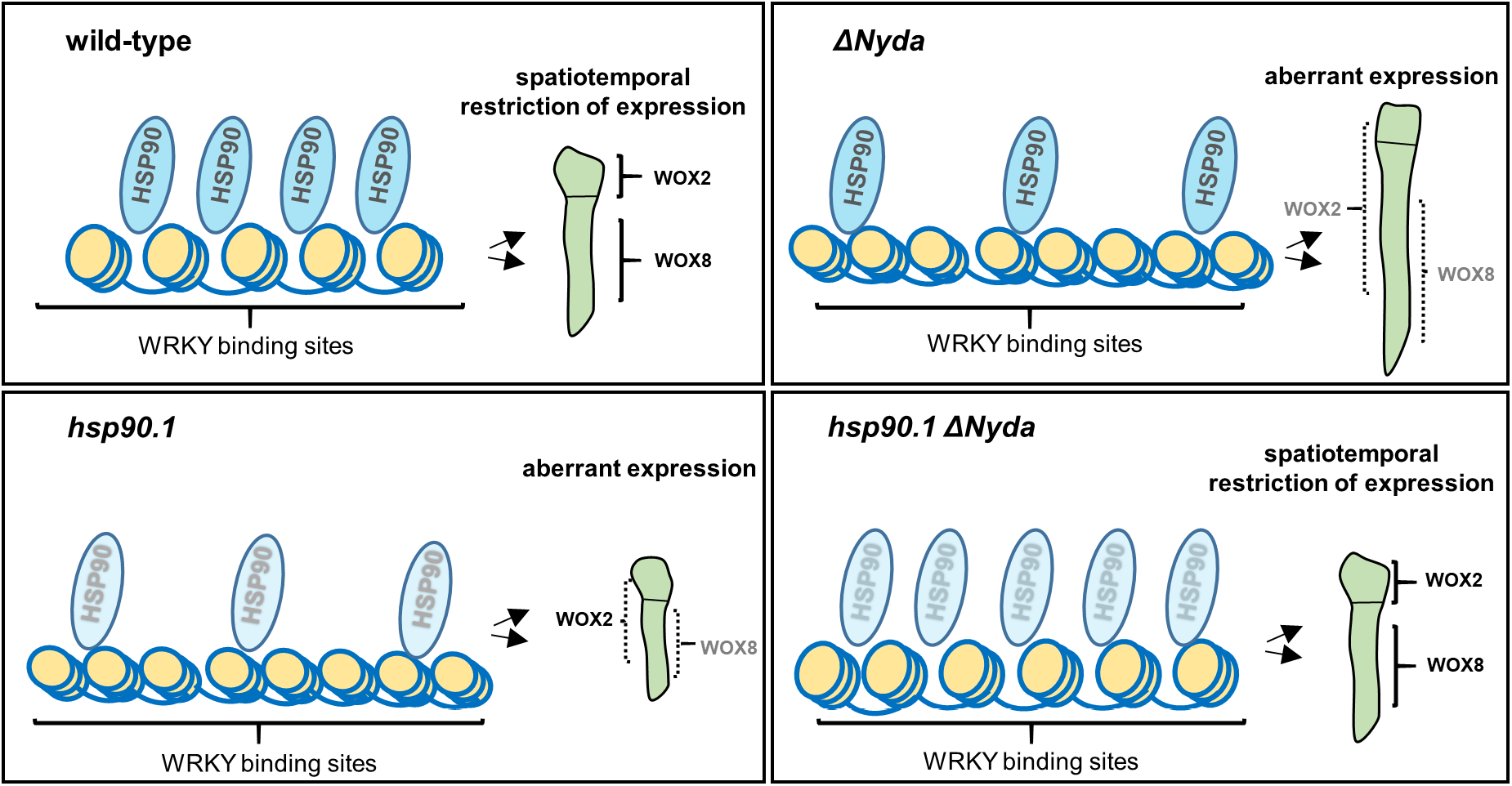
Proposed model for the interplay of HSP90s and YDA in the control of the spatiotemporal expression of *WOX8* and *WOX2* during early embryo development. HSP90s depletion or constitutive activation of YDA results in alterations in nucleosome occupancy on WRKY binding sites in the promoters of *WOX* genes. Increased deposition of nucleosomes leads to displacement of WRKY2 transcription factors contributing to lower transcriptional rates and aberrant expression of *WOX8* and *WOX2* resulting in disturbed apical and basal cell fate determination, as well as impaired growth of the proembryo in *hsp90.1* and *ΔNyda* mutants. Restoration of nucleosome distribution and HSP90 levels in *hsp90.1 ΔNyda* double mutants suppresses the severe embryo phenotype of *ΔNyda* and restores the expression patterns of *WOX* genes.

## Acknowledgements

This work has been supported by Czech Science Foundation GAČR (project No. 17-24500S) and by ERDF project “Plants as a tool for sustainable global development” (No. CZ.02.1.01/0.0/0.0/16_019/0000827). We are grateful to Prof. Thomas Laux (University of Freiburg) for kindly providing *proWOX8:NLS-YFP* and *proWOX2:NLS-YFP* lines that have been used throughout this study. We thank Prof. Polydefkis Hatzopoulos for kindly providing *proHSP90.1:HSP90.1-GFP* and *proHSP90.2:HSP90.2-GFP* lines, Prof. Juan Dong for *proYDA:YDA-GFP* marker line and Prof. Klaus Palme for *proDR5:GFP* line.

## Author contributions

D.S., T.T., T.V. and N.Z. performed experiments, evaluated and analysed data. J.Š. provided infrastructure, helped to plan some experiments, and contributed to data interpretation. D.S. wrote the manuscript with a contribution of T.V., T.T., M.O. A.P. and J.Š.

## Supplemental Data

**Supplemental Figure S1. Expression of HSP90s and YDA in developing embryos.** (A) Expression of *proHSP90.1:GUS, proHSP90.2:GUS* and *proYDA:GUS* in early and late globular stages of embryogenesis. Black arrows indicate the location of the embryo. (B) Expression of *proHSP90.1:HSP90.1-GFP*, *proHSP90.2:HSP90.2-GFP* and *proYDA:YDA-YFP* in different stages of embryo development as indicated above the pictures. Scale bars, 50 μm for (A) and 20 μm for (B), respectively.

**Supplemental Figure S2. Epistatic effects on embryo suspensor development in *hsp90 yda-1* and *hsp90 ΔNyda* double mutants.** Epistasis values were calculated for all pairwise combinations of double mutants. Epistasis values are presented as mean±SD (N≥20). The data were analyzed by ANOVA followed by Tukey’s test (ns is not statistically significant, * is significant at p<0.05, ** is significant at p <0.01, ** is significant at p <0.001). Results of the statistical analysis are in Table S8.

**Supplemental Figure S3. CDP orientation defects in *mpk3, mpk6* and *mpk6AEF* developing embryos.** Images show propidium-iodide stained embryos (shown in inverted black-white mode) at 2-to 4-cell stage and octant stage, respectively, of the indicated genotypes. Red arrowheads point to wrongly positioned cell walls. Scale bars, 10 μm.

**Supplemental Figure S4. Analysis of *proWOX8:NLS-YFP* and *proWOX2:NLS-YFP* expression in *hsp90, yda and mpk* mutants.** (A) Percentage of embryos of the indicated genotypes expressing *proWOX8:NLS-YFP* in early (up to 16-cell stage) and later stages (dermatogen) of early embryo development (N≥142 with the exception of *yda-1* and *ΔNyda*, where N≥37). (B) Percentage of embryos of the indicated genotypes expressing *proWOX2:NLS-YFP* in different developmental stages (N≥96).

**Supplemental Figure S5. Genetic depletion of HSP90s leads to alterations in *WOX8* transcriptional activation.** Analysis of *proWOX8:NLS-YFP* localization and signal distribution during early and later stages of early embryo development in *hsp90* mutants. From left to right: merged images of DIC with fluorescent YFP signals, fluorescent YFP signals, and semi-quantitative fluorescence intensity evaluation of *proWOX8:NLS-YFP* expression levels using pseudo colour-coded range from the scale, where dark blue represents zero intensity (150 in arbitrary units) and white represents maximum intensity (1000 in arbitrary units). Scale bars, 20 μm.

**Supplemental Figure S6. *WOX8* expression patterns in *mpk3-1, mpk6-2, mpk6AEF* and quantitative analysis.** (A) Analysis of *proWOX8:NLS-YFP* localization and signal distribution during early and later stages of early embryogenesis in the indicated genotypes. From left to right: merged images of DIC with fluorescent YFP signals and SCRI Renaissance 2200 staining with fluorescent YFP signals. Yellow arrows point to aberrant expression of *proWOX8:NLS-YFP* in the suspensor cells, red arrows indicate ectopic signal in the apical part. Scale bars, 10 μm. (B,C) Quantitative analysis of abnormal signal localization in the indicated genotypes (N=number of observed embryos; stars mark statistically significant increase in observed abnormal phenotypes, statistical analysis was performed by χ^2^ test and its results are in Table S9).

**Supplemental Figure S7. Impaired function of YDA signaling cascade affects *WOX8* expression in the embryo.** Analysis of *proWOX8:NLS-YFP* localization and signal distribution during early and later stage of early embryogenesis in *mpk3-1, mpk6-2, mpk6AEF, yda-1* and *ΔNyda* embryos. From left to right: merged images of DIC with fluorescent YFP signals, fluorescent YFP signals, and semi-quantitative fluorescence intensity evaluation of *proWOX8:NLS-YFP* expression levels using pseudo colour-coded range from the scale, where dark blue represents zero intensity (150 in arbitrary units) and white represents maximum intensity (1000 in arbitrary units). Red arrows indicate ectopic signal in the apical part. Scale bars, 20 μm.

**Supplemental Figure S8. *WOX2* expression analysis in developing embryos of *hsp90.1, hsp90.2* and *mpk3-1* mutants.** Microscopic analysis of *proWOX2:NLS-YFP* expression pattern in developing embryos of *hsp90.1, hsp90.2* (A) and *mpk3-1* (B) mutants. From left to right: merged images of DIC with fluorescent YFP signals, fluorescent YFP signals, and semi-quantitative fluorescence intensity evaluation of *proWOX2:NLS-YFP* expression levels using pseudo colour-coded range from the scale, where dark blue represents zero intensity (300 in arbitrary units) and white represents maximum intensity (2000 in arbitrary units). Scale bars, 20 μm.

**Supplemental Figure S9. Abnormal auxin distribution during embryonic development in *mpk3-1, mpk6-2* and *mpk6AEF* mutants.** Auxin maxima were visualized in lines expressing *proDR5:GFP*. (A) Microscopic analysis of *proDR5:GFP* signal distribution in late embryonic development (early heart and late heart stages) is shown for the indicated genotypes. From left to right: merged images of DIC with fluorescent GFP signals, fluorescent GFP signals, and semi-quantitative fluorescence intensity evaluation of *proDR5:GFP* expression levels using pseudo colour-coded range from the scale, where dark blue represents zero intensity (200 in arbitrary units) and white represents maximum intensity (2000 in arbitrary units). Scale bars, 20 μm. (B) The ratio of basal to the apical fluorescence signal intensity showing auxin distribution within the embryos (N>35; Kruskal-Wallis test, statistically significant differences compared to wild-type are shown, ns is not statistically significant, *** is significant at p<0.001). The results of statistical analysis are in Table S10. (C) Quantitative analysis of observed defects in auxin maxima localization in the indicated genotypes (N=number of observations).

**Supplemental Figure S10. Compensated auxin maxima in the embryos of *hsp90.1 yda-1* double mutant.** Auxin maxima were visualized in lines expressing *proDR5:GFP* in wild-type, *hsp90.1*, *yda-1* and *hsp90.1 yda-1* genetic backgrounds. From left to right: merged images of DIC with fluorescent GFP signals, fluorescent GFP signals, and semi-quantitative fluorescence intensity evaluation of *proDR5:GFP* expression levels using pseudo color-coded range from the scale, where dark blue represents zero intensity (200 in arbitrary units) and white represents maximum intensity (2000 in arbitrary units). Scale bars, 20 μm.

**Supplemental Figure S11. Positive and negative control for HSP90s-CHIP qPCR analysis.** (A) ChIP using H3 antibody followed by qPCR analysis of WRKY binding sites at the promoters of *LEC2, ACT2, WOX9, WOX8, WOX2* and *FUS3* in wild-type. (B) Fold enrichment of HSP90s at WRKY binding sites after HSP90-CHIP qPCR analysis in the promoters of *LEC2* and *ACT2*. Data are presented as means±SD (N≥three biological and technical triplicates; ANOVA followed by Scheffé’s test was applied for statistical evaluation; letters in the graph are shared by groups without statistically significant differences at the 0.05 probability level; results of statistical analysis are in Table S11).

## List of tables

**Table S1:** List of genotypes used in this study.

**Table S2:** List of primers used in this study.

**Table S3:** Statistical analysis for Fig.1B.

**Table S4:** Statistical analysis for Fig.1C.

**Table S5:** Statistical analysis for Fig.5B.

**Table S6:** Statistical analysis for Fig.6B.

**Table S7:** Statistical analysis for Fig.6D.

**Table S8:** Statistical analysis for Fig.S2.

**Table S9:** χ^2^ tests for Fig.S6B,C.

**Table S10:** Statistical analysis for Fig.S9B.

**Table S11:** Statistical analysis for Fig.S11B.

